# Dynamically stiffening biomaterials reveal age- and sex-specific differences in pulmonary arterial adventitial fibroblast activation

**DOI:** 10.1101/2023.05.11.540410

**Authors:** Mikala C. Mueller, Yanmei Du, Lori A. Walker, Chelsea M. Magin

## Abstract

Respiratory diseases like pulmonary arterial hypertension (PAH) frequently exhibit sexual dimorphism. Female PAH patients are more susceptible to the disease but have increased survival rates. This phenomenon is known as the estrogen paradox, and the underlying mechanisms are not fully understood. During PAH progression *in vivo*, human pulmonary arterial adventitial fibroblasts (hPAAFs) differentiate into an activated phenotype. These cells produce excess, aberrant extracellular matrix proteins that stiffen the surrounding pulmonary arterial tissues. Here, we employed dynamic poly(ethylene glycol)-alpha methacrylate (PEGαMA)-based biomaterials to study how the age and sex of human serum influenced hPAAF activation in response to microenvironmental stiffening *in vitro*. Results showed female and male cells responded differently to increases in microenvironmental stiffness and serum composition. Male hPAAFs were less activated than female cells on soft hydrogels and more responsive to increases in microenvironmental stiffness regardless of serum composition. Female hPAAF activation followed this pattern only when cultured in younger (age < 50) female serum or when older (age ≥ 50) female serum was supplemented with estradiol. Otherwise, female hPAAF activation was relatively high on both soft and stiffened hydrogels, with little difference in activation between the two conditions. Collectively, these results suggest that it may be possible to model the estrogen paradox observed in PAH *in vitro* and that it is critical for researchers to report cell sex and serum source when conducting *in vitro* experimentation.

## INTRODUCTION

Pulmonary arterial hypertension (PAH) is a chronic fibrotic disease where overactive tissue remodeling causes stiffening and narrowing of the arteries in the lung. This aberrant remodeling increases vascular resistance, causing the heart to work harder to pump blood through the lungs, eventually leading to right ventricular heart failure. One driver of this obstruction is the excessive deposition of extracellular matrix proteins (ECM) due to the persistence of activated fibroblasts or myofibroblasts in the tunica adventitia [1, 2]. Increased ECM deposition causes local increases in tissue stiffness that further promote dysregulation of fibroblast phenotype and alteration of pulmonary circulation [3]. Notably, many fibrotic diseases, including PAH, exhibit sexual dimorphisms [4], and there remains a need to specifically design *in vitro* experiments to investigate the mechanisms underlying sex differences in disease initiation and progression. Here, we present a biomaterials-based platform designed for studying sex dimorphism in fibroblastic activation initiated by changes in microenvironmental stiffness. Dynamically stiffening poly(ethylene glycol)-alpha methacrylate (PEGαMA)- based biomaterials that mimic the increases in microenvironmental stiffness from healthy (E = 1-5 kPa) to hypertensive pulmonary arterial tissues (E>10 kPa) [3] were designed to facilitate *in vitro* studies where cell culture medium containing human serum or fetal bovine serum (FBS) was employed to investigate the influence of how cell sex, serum sex, and serum age influence adventitial fibroblast activation in response to increased stiffness.

In the United States and Europe, 15 to 50 people per million have PAH, and females are four times more likely to be diagnosed with PAH than males [5, 6]. Despite the increased prevalence of PAH in females, males tend to have worse right ventricle function and survival rates. This disparity is known as the estrogen paradox. Studies have demonstrated that increased estrogen levels promote initiation of PAH while also protecting against severe disease progression [7–10]. The average age of diagnosis in females is 36 ± 15 years [5–7], an age at which most women are premenopausal and exhibit relatively high levels of circulating estradiol (30 to 400 pg/ml) [7]. Although female predominance in PAH incidence and survival is well established, the mechanisms underlying these differences are not well-studied and patients continue to receive the same treatments regardless of their sex [8].

There are three main cell types responsible for vascular remodeling, each residing in one of the three layers of the pulmonary artery wall: endothelial cells of the tunica intima, smooth muscle cells of the tunica media, and fibroblasts of the tunica adventitia. This investigation focuses on adventitial fibroblast activation in the pulmonary arterial walls because fibroblasts are responsible for ECM deposition, which significantly contributes to arterial remodeling [11]. Abnormal fibroblast activation increases ECM deposition, resulting in vessel stiffening and thickening. In PAH, vessel stiffening and thickening occurs through a buildup of collagens and matrix metalloproteinases (MMPs) and a concurrent decrease of their regulators, tissue inhibitors of metalloproteinases (TIMPs). A decrease of TIMPs allows for MMPs to stimulate more ECM remodeling [12]. This cycle leads to mechanistic downstream consequences for cells that interact with the ECM, such as increased activity in fibrotic signaling pathways. Activated fibroblasts differentiate into myofibroblasts in response to increased microenvironmental stiffness through mechanotransduction. Myofibroblast differentiation can be characterized by increased expression and organization of alpha-smooth muscle action (αSMA), cell-shape changes, differences in gene expression, and increased ECM production [11–13].

While fibroblast activation has been studied *in vitro* using a variety of cell sources and biomaterials, very few studies have reported cell sex and even fewer have included human serum in the cell culture medium. A study by Bertero et al. cultured hPAAFs on commercially available soft or stiff multi-well plates coated with collagen and showed increased expression of Col1a1, Col3a1, CTGF, LOX, YAP, and αSMA on stiff compared to soft [14]. Sex was not reported as a biological variable in this manuscript. Aguado et al. designed PEG-based biomaterials to recapitulate healthy and designed heart valve tissue and showed that female porcine and human valvular interstitial cells, the resident fibroblasts in heart valve leaflets, exhibited increased basal levels of αSMA stress fibers that rose in response to higher microenvironmental stiffnesses. Both female cell types showed increased activation overall compared to male cells [15]. Both studies used cell culture medium supplemented with FBS, an additive that contains amino acids, fatty acids, growth factors, hormones, and vitamins [16] to support cell growth *in vitro*. Although FBS is widely used, it presents various biological challenges such as unexpected cell growth characteristics, batch-to-batch variability, and undisclosed amounts of circulating sex hormones [17]. A few studies have moved to using human serum in cell culture medium as an alternative to FBS to avoid these limitations [17]. Aguado et al. also investigated the influence of patient-derived serum obtained before and after aortic valve replacement on fibroblast activation using biomaterials with elastic modulus values matching healthy cardiac tissue *in vitro*. Serum from before aortic valve replacement increased fibroblast activation while patient serum from after aortic valve replacement decreased activation. This differential activation was likely due to the increased immune response, cytokine signaling, and ECM proteins present before aortic valve replacement compared to the after aortic valve replacement serum [18]. Another study collected and analyzed male and female human serum for 174 different analytes. The results showed that, in addition to circulating sex hormones, 77 of the analytes were significantly different between male and female serum. A total of 40 analytes were elevated in female serum and 37 different analytes were elevated in male serum [19].

Research on the effects of circulating sex hormones on pulmonary artery cell types have investigated the influence of estrogens, specifically estradiol (E2), on the increased prevalence of PAH in females [7]. These studies have shown that increased estrogen levels promote development of PAH, but estrogen also protects against severe disease [7–10]. These studies are especially important because estrogen and progesterone levels change over the lifespan in female patients, with measurable decreases directly correlated to aging; both woman and men that develop PAH tend to have higher levels of circulating E2 [7].

Here, we build on this work by presenting a strategy to model pulmonary artery adventitial fibroblast activation using PEGαMA hydrogels that accurately replicate the stiffness of healthy distal pulmonary artery tissue (1-5 kPa) and can undergo a secondary photopolymerization reaction to stiffen the hydrogel (>10 kPa) to replicate vessel stiffening [3]. Female or male hPAAFs were grown on these hydrogels in cell culture media supplemented with FBS to observe the inherent differences in fibroblast activation in response to matrix stiffening, where fibroblast activation was measured by αSMA expression. Human sera pooled by sex and grouped by age (< 50 or ≥ 50 years old) were then used to supplement the cell culture medium to examine how sex and age affected fibroblast activation in response to matrix stiffening. Male hPAAFs demonstrated responses consistent with previous studies, including those where sex was not reported as a biological variable. Male hPAAF activation was relatively low on soft surfaces and significantly increased in response to dynamic stiffening independent of serum composition. Female hPAAFs exhibited higher levels of activation on soft substrates when cultured in FBS, with older female serum leading to no statistically significant increases in activation in response to microenvironmental stiffening.

Activation of female cells in response to stiffening was only different when the cells were cultured with younger female serum containing relatively high levels of estradiol or when estradiol was added to older female serum. These results highlight the importance of considering sex and serum source when designing experiments to model sexually dimorphic diseases, such as PAH, *in vitro*.

## MATERIALS AND METHODS

### PEGαMA Synthesis

Poly(ethylene glycol)-hydroxyl (PEG-OH; 8-arm, 10 kg mol^−1^; JenKem Technology) was dissolved in anhydrous tetrahydrofuran (THF; Sigma-Aldrich) in a flame-dried Schlenk flask and purged with argon. Sodium hydride (NaH; Sigma-Aldrich) was added with a funnel into the reaction at 3x molar excess to PEG-hydroxyl groups and reacted for 45 minutes at room temperature. Ethyl 2-(bromomethyl)acrylate (EBrMA; Ambeed, Inc.) was added drop-wise at a 6x molar ratio to PEG-OH groups, and the reaction was stirred at room temperature for 48 hours protected from light. The mixture was neutralized with 1 N acetic acid until gas evolution ceased and filtered through Celite 545 (EMD Millipore). The solution was concentrated by rotary evaporation at 50 °C and precipitated in cold diethyl ether (Fisher Scientific) at 4 °C overnight. The solid product was then dried under vacuum overnight at room temperature. The product was purified using dialysis against deionized water (1 kg mol^−1^ MWCO, ThermoFisher) for four days, then flash frozen at −80 °C, and lyophilized to give the final product. The functionalization of the product was verified by 1H NMR. 1H NMR (300 MHz, CDCl3): d (ppm) 1.36 (t, 3H, CH3–), 3.71 (s, 114H, PEG CH2-CH2), 4.29 (t, s, 4H, –CH2–C(O)–O–O, –O–CH2–C(=CH2)–), 5.93 (q, 1H, –C=CH2), 6.34 (q, 1H,–C=CH2) [20]. PEGαMA with functionalization over 90%, as measured by comparing the αMA vinyl end group peak to that of the PEG backbone, was used in subsequent experiments (Figure S1).

### Hydrogel Formation

PEGαMA hydrogels were formed using a previously published protocol, where a design of experiments approach selected the best formulation [21]. Briefly, PEGαMA hydrogels were made by dissolving the PEGαMA in pH 8.4 4-(2-hydroxyethyl)-1-piperazineethanesulfonic acid (HEPES; Life Technologies) at 12.5-15.5 weight percent (wt%). 1-4,Dithiothreitol (DTT; Sigma-Aldrich) and KCGPQGIWGQCK (MMP-2 degradable peptide; GenScript) were used as crosslinkers in a 70:30 molar ratio respectively and were dissolved in 15 mM tris(2-carboxyethyl)phosphine hydrochloride (TCEP; Sigma-Aldrich). Fibronectin mimetic peptide (CGRGDS; GL Biochem) was used at a 2 mM concentration to facilitate cell adhesion to hydrogels and was dissolved in TCEP.

Hydrogels for rheological evaluation were formed by placing 40-µL drops of precursor solution between two glass slides covered in parafilm and allowing 30 min for polymerization. Hydrogels for stiffening experiments were swelled in phosphate-buffered saline (PBS; HyClone) with 2.2 mM lithium phenyl-2,4,6-trimethylbenzoylphosphinate (LAP; Sigma-Aldrich) photoinitiator overnight. The hydrogels were stiffened by exposure to ultraviolet (UV) light at 365 nm, 10 mW cm^-2^ (Omnicure, Lumen Dynamics) for 5 min. Previous studies have shown no significant differences in fibroblast viability and transcriptome when the cells were exposed to 365 nm light [22].

Hydrogels for cell culture experiments were fabricated by placing 90-µL drops of precursor solution between a 3-(trimethoxysilyl)propyl acrylate (Sigma-Aldrich) silanated 18-mm glass coverslip and a glass slide treated with SigmaCote (Sigma-Aldrich), then allowing 30 min for polymerization. These hydrogels were soaked in growth media (basal medium with 0.1% insulin, 0.1% gentamicin sulfate – amphotericin, 0.1% human epidermal growth factor, 0.2% human fibroblast growth factor-beta, and 5% FBS; Lonza) overnight at 37 °C prior to cell seeding.

### Hydrogel Characterization

Rheology was performed as previously described [23]. Mechanical properties of soft and stiffened hydrogels were assessed on a Discovery HR2 rheometer (TA Instruments) using an 8-mm parallel plate geometry and the Peltier plate set at 37 °C. The geometry was lowered until 0.03 N axial force was read and the gap distance was recorded. The gap distance was adjusted starting at 25% compression until the storage modulus (G’) stabilizes under compression, with the percent compression at which G’ plateaued used for measurement samples [24]. The samples were exposed to frequency oscillatory strain with a frequency range of 0.1 to 100 rad s^−1^ at 1% strain.

The elastic modulus (E) was calculated using rubber elastic theory, assuming a Poisson’s ratio of 0.5 for bulk measurements of elastic hydrogel polymer networks.

### Sex-Hormone Analysis

Human serum from healthy donors (Innovative Research) was pooled together in equal parts by age (Table 1).

**Table 1.** Human serum information.

| Sex | Age | Pooled |
| --- | --- | --- |
| Female | 63 | Age ≥ 50 |
| Female | 50 |  |
| Female | 43 | Age < 50 |
| Female | 36 |  |
| Female | 29 |  |
| Male | 64 | Age ≥ 50 |
| Male | 56 |  |
| Male | 47 | Age < 50 |
| Male | 20 |  |
| Male | 20 |  |

Testosterone concentrations in serum were measured with an enzyme-linked immunosorbent assay (ELISA; Abcam ab174569) following the manufacturer’s protocol. Estradiol and Progesterone concentrations in serum were measured by the Clinical and Translational Research Center at the University of Colorado, Anschutz Medical Campus using a chemiluminescent immunoassay (Beckman Coulter) following the manufacturer’s protocol.

### Sex-Specific hPAAF Activation

#### Investigation of Commercially Available Cell Lines

Female (Creative Bioarray (donor age = 11) and male AccelGen (donor age = 2) hPAAFs were expanded at 37°C and 5% CO_2_ on tissue-culture-treated polystyrene in growth media (basal medium with 0.1% insulin, 0.1% gentamicin sulfate – amphotericin, 0.1% human epidermal growth factor, 0.2% human fibroblast growth factor-beta, and 5% FBS; Lonza). Female and male hPAAFs were seeded at 20,000 cells cm^-2^ onto the hydrogels in growth media and used between passage 3-6. The next day the media was changed to activation media (growth media with 1% serum). Serum used in these experiments included FBS (Cytiva) or human serum pooled by sex and age. All hPAAFs were cultured on soft hydrogels for 7 days with 2.2 mM LAP added to the activation media on day 6. Half of the hydrogels were exposed to UV light at 365 nm, 10 mW cm^-2^ (Omnicure, Lumen Dynamics) for 5 min on day 7 to stiffen these samples, then media was changed to ensure the LAP was rinsed out. Soft hydrogels were collected for analysis on day 7 and stiffened hydrogels were collected on day 9, as previously described [23].

Samples were fixed with 4% (vol/vol) paraformaldehyde (PFA, Electron Microscopy Sciences) and permeabilized with 0.5% Triton X-100 (Thermo Scientific) solution for 20 minutes, quenched with 100 mM glycine (Fisher Scientific) for 15 minutes, and blocked with 5% bovine serum albumin (BSA; Sigma-Aldrich) for 1 hour. All steps for immunostaining were performed at room temperature. Primary alpha smooth muscle actin (αSMA) antibody (mouse anti-human αSMA monoclonal antibody (MA5-11547); Invitrogen) was diluted 1:250 in 3% BSA and 0.1% Tween 20 (Thermo Scientific) in PBS (IF solution), then incubated for 1 hour. Samples were washed three times with IF solution. Goat anti-mouse Alexa Fluor-555 (Invitrogen) secondary antibody was diluted 1:250 in IF solution and Actin 488 ReadyProbes (AlexaFluor 488 phalloidin; Invitrogen) were added to the solution according to the manufacturer’s instructions. Samples were incubated with this solution for 1 hour and washed three times with IF solution. 4′,6-diamidino-2-phenylindole (DAPI; BioLegend) was diluted 1:10,000 from a 5 mg mL^-1^ stock solution in deionized water, incubated for 20 minutes, and washed once with deionized water, once in PBS, and once in IF solution. Samples were mounted using Prolong Gold Antifade (Invitrogen) for imaging and sealed with nail polish (Electron Microscopy Sciences) for storage.

Six images were taken at random fields of view on each sample using an Olympus upright epifluorescence microscope (BX-63) and then processed with ImageJ software (NIH). Fibroblast activation was defined as the percentage of cells with αSMA present over the total number of cells counted by DAPI-stained nuclei ((cells positive for αSMA/total # cells) *100%).

#### Validation in Female and Male Donor Cells

Female and male hPAAFs were purchased from the Pulmonary Hypertension Breakthrough Initiative (PHBI) to verify the differences observed in the first cell lines across multiple donors at ages close to the average age of PAH diagnosis (Table 2). Activation experiments were run using the same procedures described above using 1% age < 50 female or male serum, age ≥ 50 female or male serum, or age ≥ 50 female serum supplemented with additional estradiol. Estradiol (Innovative Research of America) was dissolved in 100% ethanol to a concentration of 1 ng/mL. Next, 30 µL of 1 ng/mL estradiol in ethanol was added to 50 mL of media containing 1% older female serum to adjust the final estradiol concentration to match the average younger female serum (115.7 pg/mL). Vehicles were samples treated with media made with 30 µL of ethanol added to 50 mL of media.

**Table 2.** PHBI hPAAF donor information.

| PHBI App # | Sex | Age |
| --- | --- | --- |
| PHBI-AH-014 | Female | 28 |
| PHBI-BA-054 | Female | 33 |
| PHBI-UA-015 | Female | 36 |
| PHBI-BA-033 | Male | 24 |
| PHBI-UA-008 | Male | 25 |
| PHBI-AH-011 | Male | 30 |

### Statistical Methods

Data were assessed for normality using Anderson-Darling and Shapiro-Wilk tests. A Brown-Forsythe test determined equal sample variance between PHBI donor results, which were pooled for further analysis. After normality was confirmed and data were pooled, a two-way analysis of variance (ANOVA) with Tukey’s honest statistical difference (HSD) test for multiple comparisons was performed. Values were reported as mean ± standard deviation (SD) and considered significantly different if the p-value < 0.05, with each data point representing a different technical replicate (three technical replicates per biological replicate and three biological replicates per condition). Statistical analysis was performed using GraphPad Prism software.

A custom design of experiments generated with JMP software (SAS, Cary, NC) analyzed the impact of each variable on fibroblast activation. The level of each variable (e.g., stiffness, cell sex, serum sex, and serum age) and the associated hPAAF activation results were entered into the custom design and analyzed with a least-squares regression model to plot response curves and determine the most significant factors.

## RESULTS

### Hydrogel Synthesis and Characterization

Soft hydrogels were designed to mimic the stiffness of healthy pulmonary artery tissue and stiffened into the range of hypertensive tissue. Soft hydrogels were polymerized by an off-stoichiometry (0.375 thiols to αMA moieties) Michael addition reaction under slightly basic conditions with two crosslinkers (DTT and an MMP2-degradable crosslinker), along with a cell adhesion peptide (CGRGDS), which mimics binding sites on the ECM protein fibronectin (Figure 1A). Stiffened hydrogels were formed by soaking soft hydrogels in a photoinitiator (LAP) overnight and then applying UV light to the hydrogels to initiate homopolymerization between the unreacted αMA moieties (Figure 1A). The soft hydrogels achieved an average elastic modulus value of 4.6 ± 1.0 kPa, matching the range for healthy pulmonary arteries (Figure 1B). Likewise, an elastic modulus of 19.7 ± 1.3 kPa was measured for the stiffened hydrogels, matching the range for hypertensive arterial tissue (Figure 1B).

**Figure 1.**
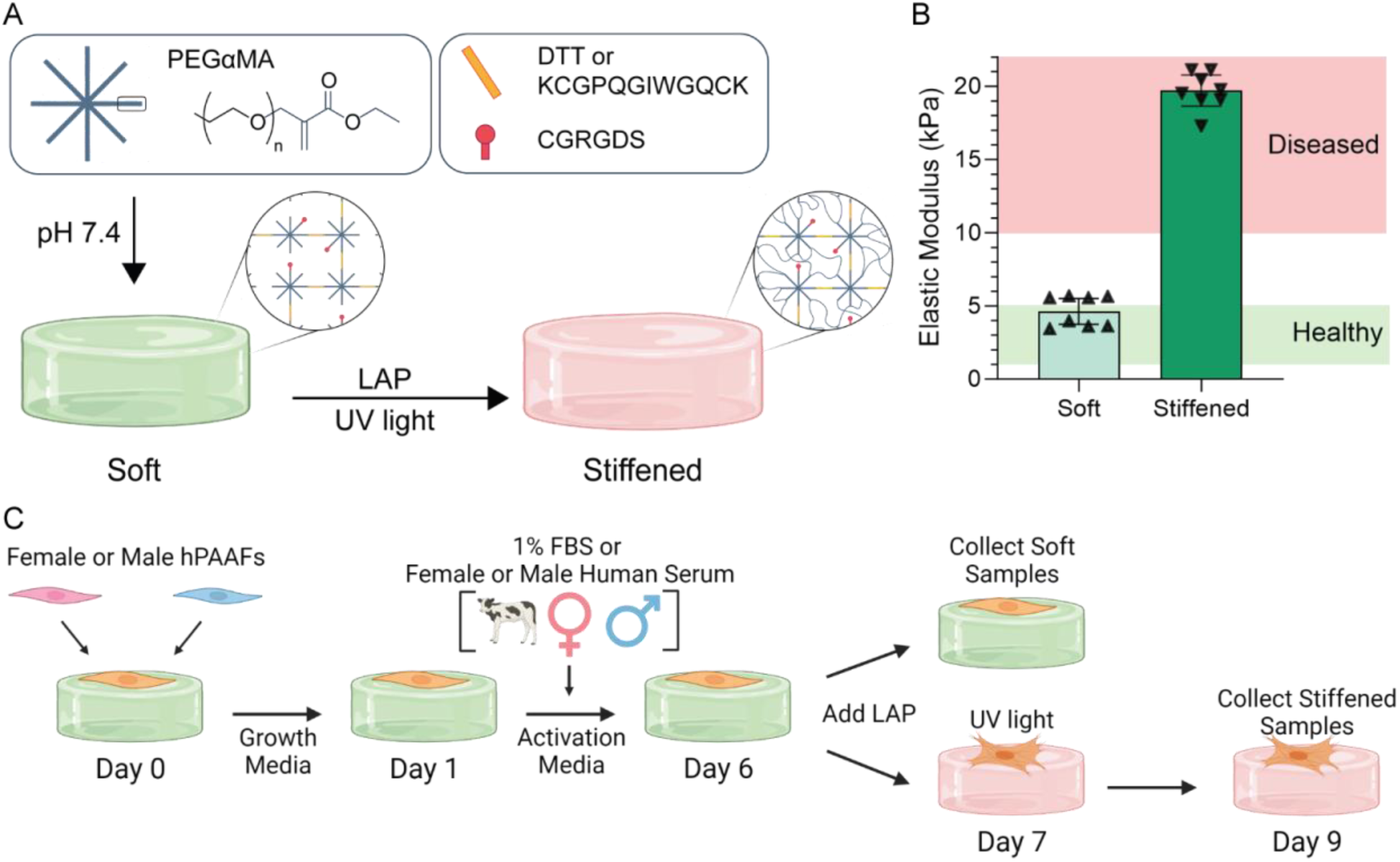
Characterization of PEGαMA hydrogels. A) Schematic of the hydrogel formulation containing PEGαMA, DTT, MMP-2 degradable crosslinker, and CGRGDS. Hydrogel components were reacted under basic conditions to form soft hydrogels. Incubation in LAP and exposure to UV light initiated a homopolymerization reaction to stiffen the hydrogels. B) Rheological measurements showed the average elastic modulus (E) of soft hydrogels (E = 4.63 ± 1.0 kPa) was within the range of healthy pulmonary arteries (1-5 kPa) and the average elastic modulus of stiffened hydrogels (E = 19.70 ± 1.3 kPa) increased into the range of pathologic pulmonary arteries (>10 kPa). Columns represent mean ± SD, n = 8. C) Schematic representation of the timeline used for cell experiments and created with BioRender.com.

Female and male hPAAFs were cultured on soft hydrogels for 7 days, then half of the samples were stiffened. Soft samples were collected and analyzed on day 7, while stiffened samples were collected for analysis on day 9. Cells collected from soft hydrogels on days 7 and 9 exhibited baseline activation levels that were not statistically different (Figure S2). Cells were seeded on hydrogels in complete growth medium containing 5% FBS. On day 1 media was changed to activation media containing 1% FBS or sex- and age-specific human serum. On day 6 the LAP photoinitiator was added to the cell culture medium for all samples and allowed to soak into the hydrogels overnight. The next day (day 7), UV light was applied to the samples, stiffening hydrogels for the stiffened condition. Media was changed the next day to remove any unreacted photoinitiator and the cells were grown for two more days on the stiffened hydrogel before the samples were collected and analyzed (Figure 1C).

### Sex-Specific hPAAF Activation in Response to Increases in Microenvironmental Stiffness

Female or male cells were cultured in a medium containing 1% FBS on soft or stiffened hydrogel conditions to measure the differences in hPAAF activation related to cell origin in response to increases in microenvironment stiffness. Samples were immunostained for αSMA, a myofibroblast marker, to measure the percent of activated hPAAFs on the two hydrogel conditions after the experiment. F-actin and nuclei (DAPI) were also stained to visualize cell morphology and quantify the total number of cells, respectively (Figure 2A and B). Female hPAAFs were 62.6% ± 20.5% activated on soft hydrogels and 67.9% ± 20.6% activated on stiffened hydrogels (Figure 2C). Male hPAAFs were 44.6% ± 13.6% activated on soft hydrogels and 81.8% ± 9.4% activated on stiffened hydrogels (Figure 2C). Quantification of αSMA expression by image analysis was verified by western blot (Figure S3). Female hPAAFs demonstrated a non-significant 1.08-fold change in activation on stiffened substrates compared to soft, while male hPAAFs showed a significant 1.83-fold increase in activation.

**Figure 2.**
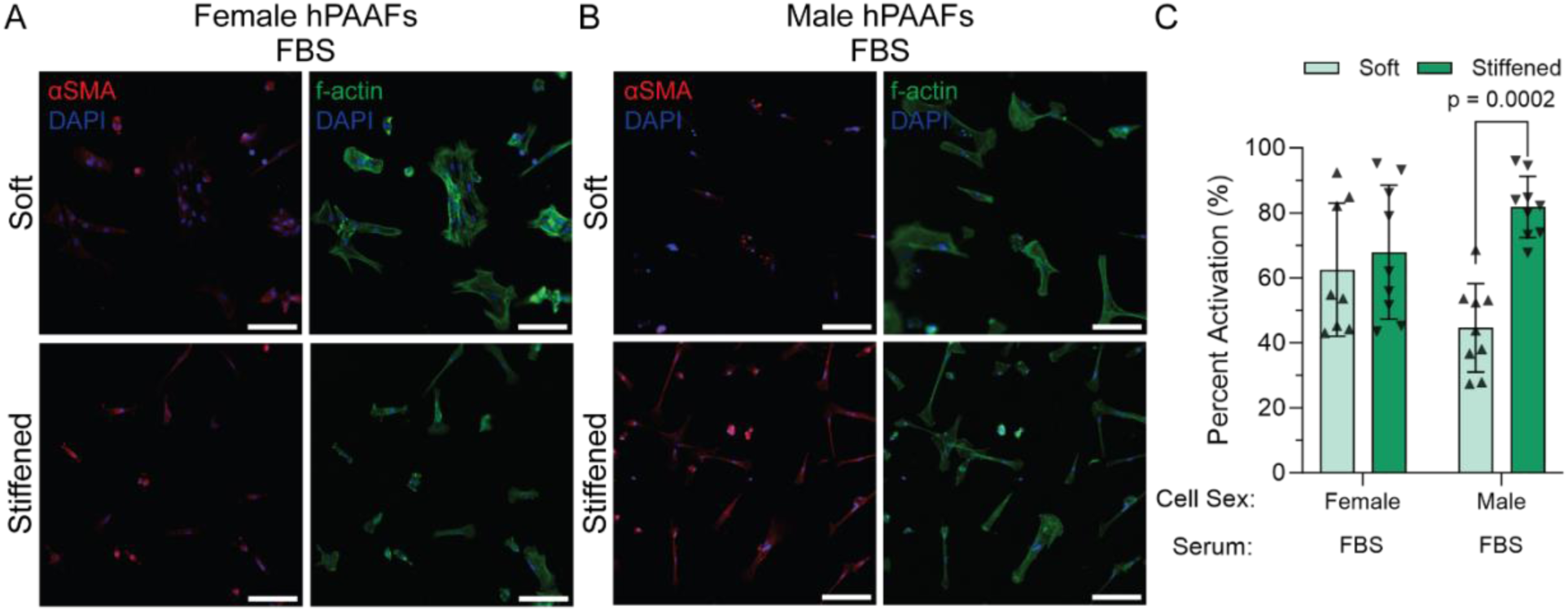
Female and male hPAAF activation in response to increases in microenvironmental stiffness. Representative fluorescence microscopy images of A) female and B) male hPAAFs grown on soft and stiffened conditions in FBS were stained for αSMA (red), f-actin (green), and DAPI (blue). C) Quantification of the percent of activated fibroblasts that showed no significant differences in female hPAAF activation on stiffened hydrogels compared to soft controls. A significant increase in male hPAAF activation was measured (*p* = 0.0002), indicating that female cells grown in FBS were not as sensitive to increases in microenvironmental stiffness. Columns represent mean ± SD, n = 3, two-way ANOVA, Tukey HSD. Scale bar: 100 µm.

### Concentration of Sex Hormones in Human Serum Samples

The concentrations of the sex hormones estradiol, progesterone, and testosterone were analyzed in serum from each donor (Table 3). The concentration of progesterone was comparable between both age ≥ 50 and age < 50 female serum, and with age ≥ 50 male serum. Age < 50 male serum had higher levels of progesterone present than the other human serum, while FBS has the highest concentration of progesterone. Estradiol concentrations were similar in both age ≥ 50 and age < 50 male serum, while estradiol was the highest in age < 50 female serum followed by age ≥ 50 female serum. FBS had the lowest concentration of estradiol present. Testosterone was comparable between FBS, age ≥ 50 and age < 50 female serum. Age ≥ 50 and age < 50 male serum had similar levels of testosterone, but more testosterone than the female serum or FBS.

**Table 3.**
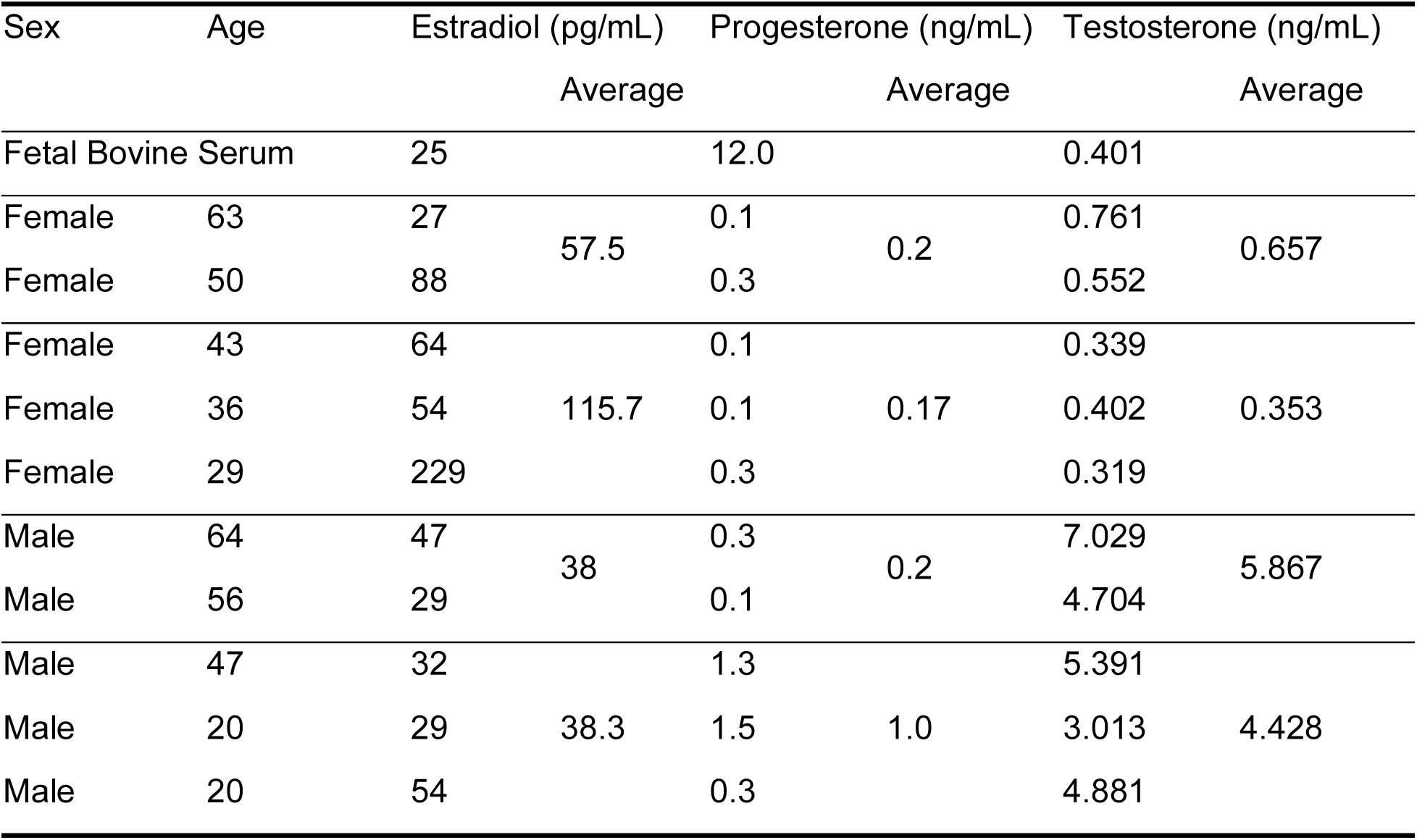
Concentrations of estradiol, progesterone, and testosterone in each serum used.

### Sex-Matched Human Serum and Microenvironmental Stiffness Influence Activation in Commercially Available hPAAF Cell Lines

hPAAFs were cultured in a medium supplemented with 1% sex-matched human serum, pooled by age, on soft and stiffened hydrogels to investigate the effects of sex- and age-related changes in serum composition and circulating sex-hormone concentration on hPAAF activation in response to increases in microenvironmental stiffness. Serum from individual donors was pooled into four different groups separated by sex and age (Table 1). Cells were cultured in younger (age < 50) or older (age ≥ 50) sex-matched serum and samples were immunostained for αSMA, f-actin, and DAPI (Figure 3A-E). Female hPAAFs grown on soft conditions were 50.5% ± 7.5% activated when grown in younger female serum and 72.9% ± 9.4% on stiffened conditions representing a 1.44-fold change in activation. Male hPAAFs demonstrated an average activation of 45.1% ± 5.8% on soft conditions and 64.6% ± 12.5% on stiffened conditions when grown in younger male serum, representing a fold change in activation of 1.43 (Figure 3C). When grown in older female serum, female hPAAF activation was 64.3% ± 9.6% on soft conditions and 55.1% ± 12.2% on stiffened conditions and had a fold change in activation of 0.86. In contrast, when grown in older male serum, male hPAAF activation was 62.2% ± 7.3% on soft conditions and 86.7% ± 7.4% on stiffened conditions, with a fold change in activation of 1.39. Over 60% of female and male cells cultured on soft hydrogels in sex-matched, older serum were activated, and there was no statistically significant increase in female cell activation upon stiffening. However, the activation for male cells was significantly higher on stiffened surfaces compared to female cell activation, indicating that circulating sex hormone concentrations may influence female cell activation levels.

**Figure 3.**
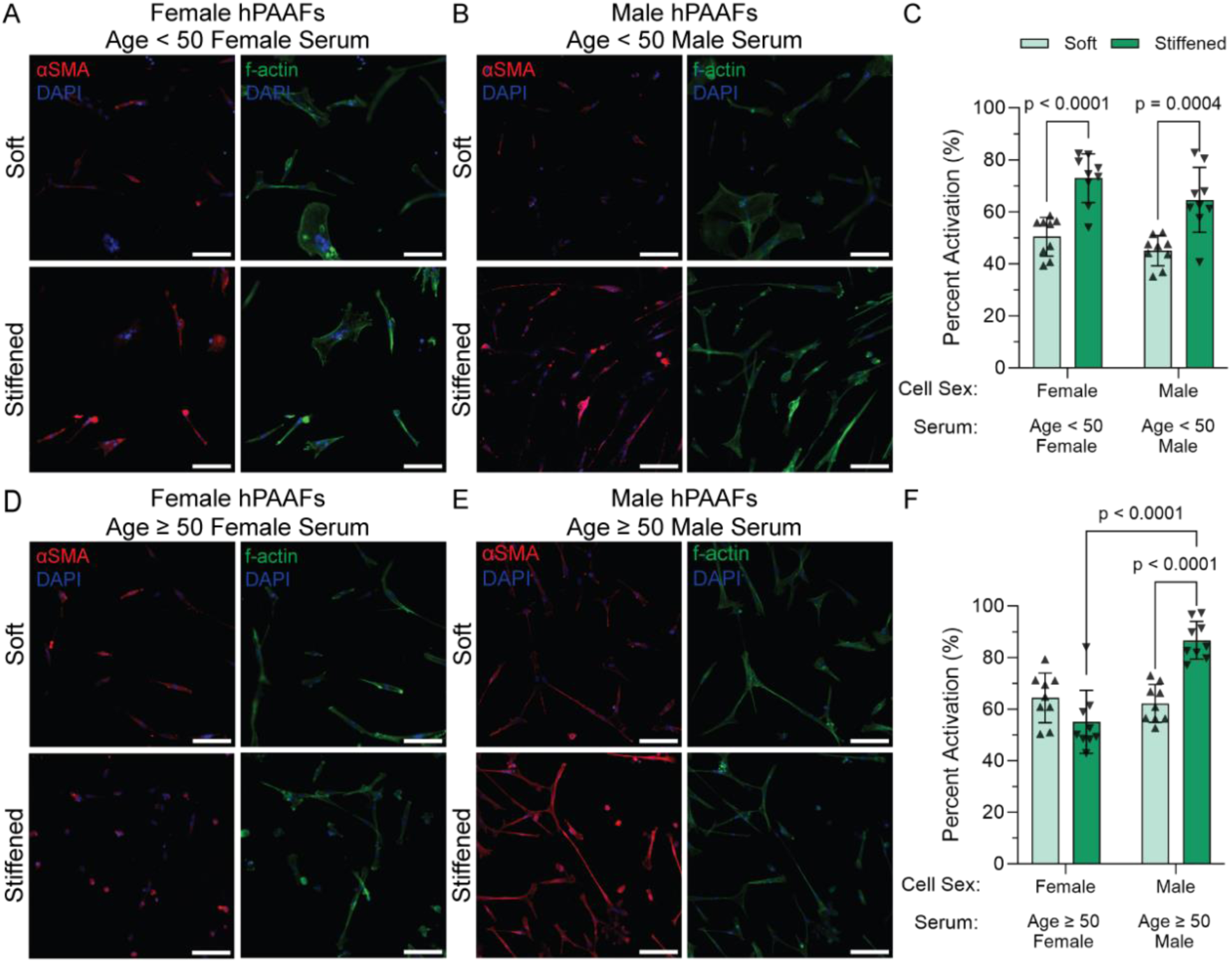
Female and male hPAAF activation in sex-matched human serum. Representative fluorescence microscopy images of A) female hPAAFs in age < 50 female serum and B) male hPAAFs grown in age < 50 male serum on soft and stiffened conditions were stained for αSMA (red), f-actin (green), and DAPI (blue). C) Quantification of the percent of activated fibroblasts of female and male hPAAFs grown in age < 50 female or male human serum, respectively, showed a significant increase in female and male hPAAF activation (*p* = <0.0001 and 0.0004, respectively) in age < 50 serum on stiffened hydrogels compared to soft controls. Similar increases in activation were observed in female and male hPAAF activation in age < 50 serum. Columns represent mean ± SD, n = 3, two-way ANOVA, Tukey HSD. Scale bar: 100 µm. Representative fluorescence microscopy images of D) female hPAAFs in age ≥ 50 female serum and E) male hPAAFs grown in age ≥ 50 male serum on soft and stiffened conditions were stained for αSMA (red), f-actin (green), and DAPI (blue). F) Quantification of the percent of activated fibroblasts of female and male hPAAFs grown in age ≥ 50 female or male human serum, respectively, showed only a significant increase in male hPAAF activation in age ≥ 50 serum was measured (*p* = <0.0001) on stiffened hydrogels compared to soft controls. There was not a significant increase in activation of the female cells grown in age ≥ 50 female serum. Columns represent mean ± SD, n = 3, two-way ANOVA, Tukey HSD. Scale bar: 100 µm

### Sex-Mismatched Human Serum and Microenvironmental Stiffness Influence Activation in Commercially Available hPAAF Cell Lines

To further explore the interplay between cell sex and circulating sex hormones, 1% human serum was added to the cell culture media for cells of the opposite sex. Specifically, male serum was added to female cells and female serum was added to male cells. Samples were immunostained for αSMA, f-actin, and DAPI (Figure 4A-E). Male hPAAF activation in age < 50 female serum on the soft condition was 50.3% ± 7.8% and 78.9% ± 6.0% on the stiffened condition, while the female hPAAF activation in age < 50 male serum was 76.2% ± 13.6% on the soft condition and 88.0% ± 7.6% on the stiffened condition (Figure 4C). Male hPAAFs in female serum showed a 1.57-fold change in activation, while female hPAAFs in male serum resulted in a 1.15-fold change in activation. Female hPAAF activation when grown in age ≥ 50 male serum was 75.4% ± 5.7% on soft conditions and 81.6% ± 8.3% on stiffened conditions with a fold change in activation of 1.08, whereas the activation of male hPAAFs grown in age ≥ 50 female serum was 87.8% ± 9.7% on soft conditions and 94.7% ± 5.0% on stiffened conditions with a fold change in activation of 1.08 (Figure 4F). Female and male cells cultured in sex mis-matched serum age < 50 showed similar trends to cells cultured in FBS, while cells cultured in age ≥ 50 sex mis-matched serum all showed higher activation levels.

**Figure 4.**
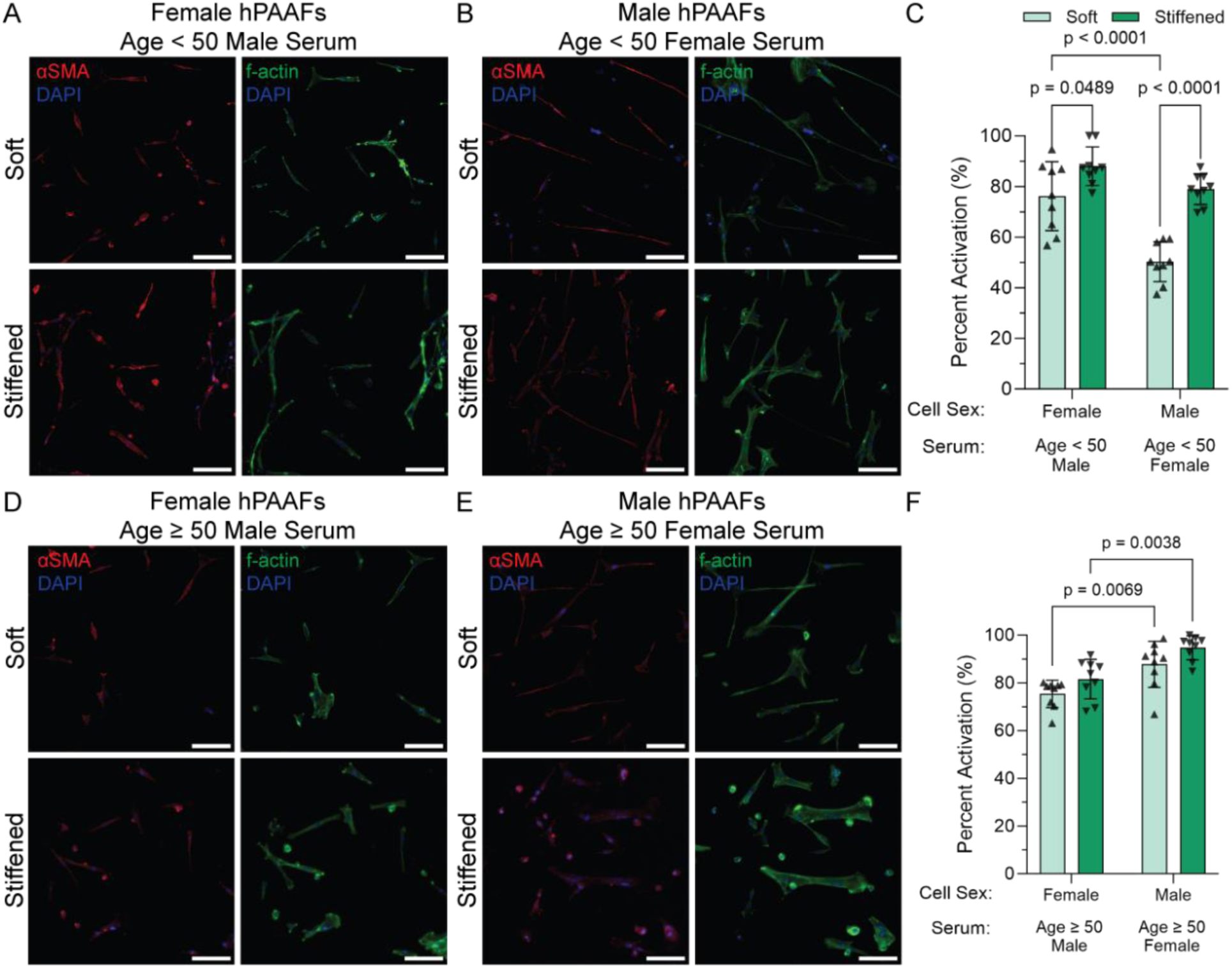
Female and male hPAAF activation in sex-mis-matched human serum. Representative fluorescence images of A) male hPAAFs in age < 50 female serum and B) female hPAAFs age < 50 male serum grown on soft and stiffened conditions were stained for αSMA (red), f-actin (green), and DAPI (blue). C) Quantification of activated fibroblasts showed significant increases (*p* = <0.0001 and 0.0489, respectively) in both male and female hPAAF activation in sex-mismatched serum on stiffened hydrogels compared to soft controls, though the difference in activation of female hPAAFs in age < 50 male serum was minimal. A significant increase in activation was measured (*p* = <0.0001) in female hPAAFs on soft hydrogels compared to male hPAAFs on soft hydrogels. Columns represent mean ± SD, n = 3, two-way ANOVA, Tukey HSD. Scale bar: 100 µm. Representative fluorescence microscopy images of D) female hPAAFs in age ≥ 50 male serum and E) male hPAAFs grown in age ≥ 50 female serum on soft and stiffened conditions were stained for αSMA (red), f-actin (green), and DAPI (blue). F) Quantification of the percent of activated fibroblasts of female and male hPAAFs grown in age ≥ 50 male or female human serum, respectively, showed no significant increase in activation of either female or male hPAAFs in sex-mis-matched age ≥ 50 serum on stiffened hydrogels compared to soft controls. The activation seen in female hPAAFs was significantly lower in age ≥ 50 male serum than male hPAAFs were in age ≥ 50 female serum on both soft and stiffened conditions. Columns represent mean ± SD, n = 3, two-way ANOVA, Tukey HSD. Scale bar: 100 µm.

There was a larger difference in activation between soft and stiffened conditions in younger serum than in older serum, regardless of sex. We further investigated these differences by evaluating serum pooled by sex and age using a human cytokine antibody array with 42 targets (Figure S4). Results showed increases in some biochemical cues that could increase cellular activation, such as interleukin 1β (IL-1β), chemokine (C-X-C motif) ligand 1 (CXCL1), monocyte chemoattractant protein-1 (MCP-1), CC chemokine ligand 17 (CCL17), CCL5, platelet-derived growth factor-BB (PDGF-BB), and transforming growth factor β-1 (TGFβ-1) in older male serum. Younger male serum exhibited increased levels of IL-15 and IL-13. No increases in the 42 cytokines evaluated here were detected in older female serum compared to young. In fact, younger female serum exhibited increases in IL-3, CXCL-1, CXCL-12, and PDGF-BB compared to older (Figure S4).

### Sex-Matched Human Serum and Microenvironmental Stiffness Modulate Activation in Male and Female Donor Cells

Male hPAAFs from three donors acquired from the PHBI were cultured with human male serum to verify differences in activation related to microenvironmental stiffening between younger and older serum composition observed in commercially available cell lines. Cultures were supplemented with younger (age < 50) male serum or older (age ≥ 50) male serum, and samples were immunostained for αSMA, f-actin, and DAPI. Male PHBI donor fibroblasts supplemented with age < 50 male serum were 42.3% ± 8.8% activated when grown on soft hydrogels and 57.8% ± 5.6% on stiffened hydrogels with a 1.37-fold change in activation (Figure 5A). Activation when supplemented with age ≥ 50 male serum was increased on both substrates with 68.9% ± 8.3% on soft conditions and 82.9% ± 5.8% on stiffened conditions with a fold change in activation of 1.21 (Figure 5B).

**Figure 5.**
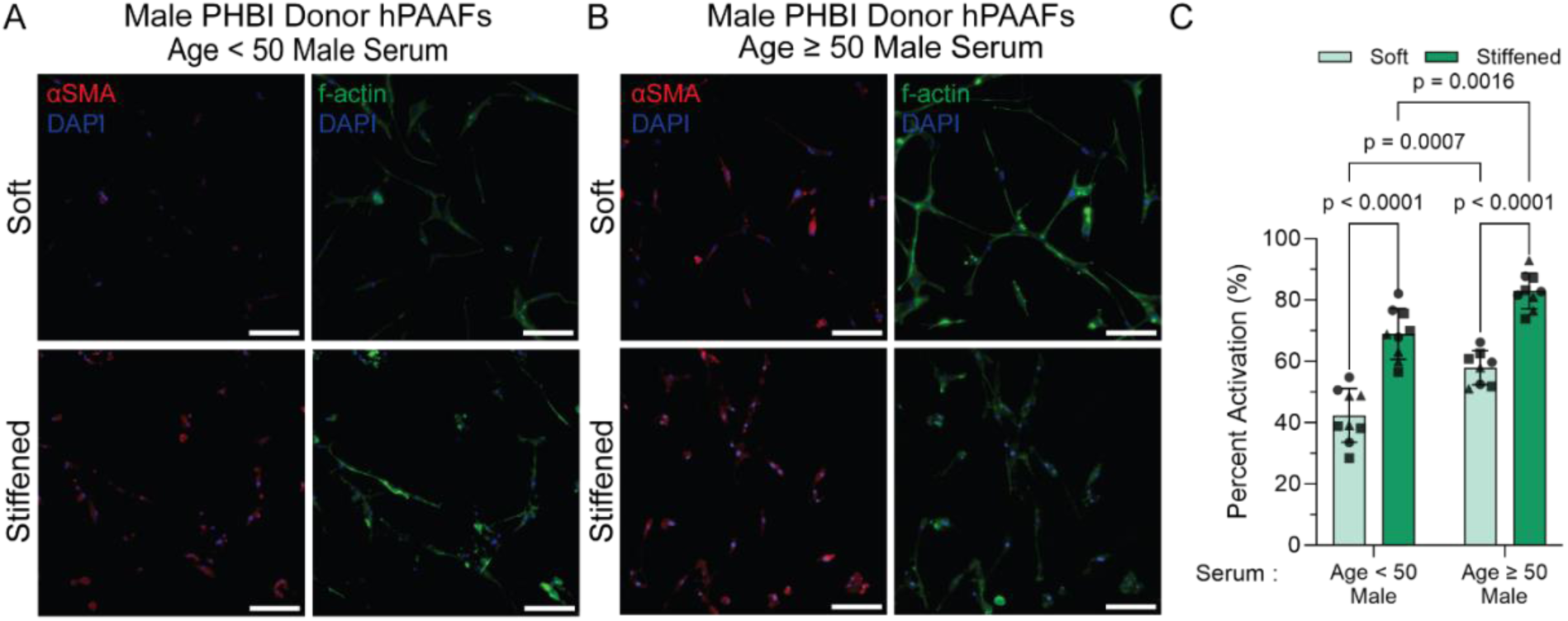
Male PHBI donor hPAAF activation in male human serum. Representative fluorescence microscopy images of A) male PHBI donor hPAAFs in age < 50 male serum and B) age ≥ 50 male serum on soft and stiffened conditions, stained for αSMA (red), f-actin (green), and DAPI (blue). Representative images are from one donor. C) Quantification of activated male PHBI donor hPAAFs cultured in age < 50 and age ≥ 50 male human serum. Each donor is represented with a different symbol on the graph. hPAAF activation significantly increased (p < 0.0001) in age < 50 male serum on stiffened hydrogels compared to soft controls. Male hPAAFs in age ≥ 50 male serum had a significant increase in activation (p < 0.0001) on stiffened samples compared to soft. A significant increase in activation was observed in hPAAFs supplemented with age ≥ 50 male serum compared to age < 50 male serum on soft samples (p = 0.0007) and stiffened samples (p = 0.0016). Columns represent mean ± SD, n = 3, and each symbol represents a different donor, two-way ANOVA, Tukey HSD. Scale bar: 100 µm.

Female hPAAFs from three donors acquired from PHBI were cultured with female human serum to verify that differences in activation related to microenvironmental stiffening between younger and older serum composition were highly influenced by estradiol concentration. Cells from each donor were expanded and seeded onto hydrogels as previously described. Cultures were supplemented with younger (age < 50) female serum, older (age ≥ 50) female serum, or older (age ≥ 50) female serum plus estradiol. Samples were immunostained for αSMA, f-actin, and DAPI (Figure 6A-C). Female PHBI donor fibroblast activation in age < 50 female serum was 52.6% ± 5.3% on soft hydrogels and 78.7% ± 10.8% on stiffened hydrogels exhibiting a 1.50-fold increase in activation. When female PHBI donor fibroblasts were cultured in age ≥ 50 female serum, 63.8% ± 17.8% activation was measured on soft conditions compared to 62.5% ± 12.6% activation on stiffened conditions, resulting in a 0.98-fold increase in activation. Female PHBI donor fibroblasts cultured in the age ≥ 50 female serum with estradiol (116 pg/mL) added to match the concentration in pooled younger female serum exhibited 52.7% ± 5.4% activation on soft hydrogels with a 1.36-fold increase in activation to 71.7% ± 6.2% activation on stiffened samples. Samples treated with a vehicle control exhibited similar responses to age ≥ 50 female serum without estradiol (Figure S5).

**Figure 6.**
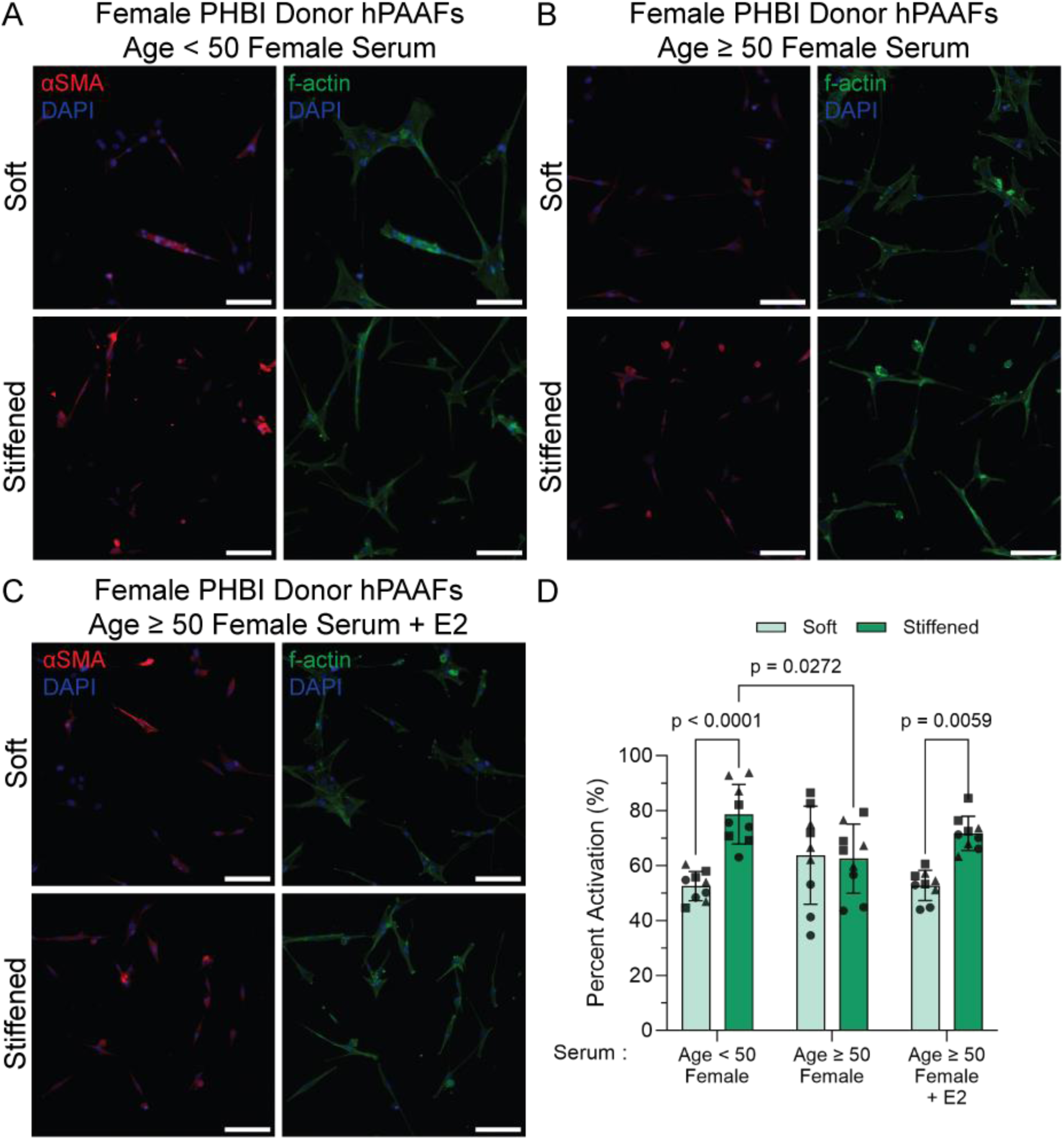
Female PHBI donor hPAAF activation in female human serum. Representative fluorescence microscopy images of A) female donor hPAAFs in age < 50 female serum, B) age ≥ 50 female serum, and C) age ≥ 50 female serum plus estradiol on soft and stiffened conditions, stained for αSMA (red), f-actin (green), and DAPI (blue). Representative images are from one donor. D) Quantification of activated female donor hPAAFs cultured in age < 50, age ≥ 50, and age ≥ 50 female human serum plus estradiol, respectively. Each symbol on the graph represents a different donor. hPAAF activation significantly increased (p = 0.0004) in age < 50 female serum on stiffened hydrogels compared to soft controls, whereas there was little change in activation between soft and stiffened conditions in age ≥ 50 female serum. When estradiol (116 pg/mL) was added to age ≥ 50 female serum, a significant increase in activation (p = 0.0059) was observed. Columns represent mean ± SD, n = 3, each symbol represents a different donor, two-way ANOVA, Tukey HSD. Scale bar: 100 µm.

### Microenvironmental Stiffness and Circulating Sex Hormone Concentrations Interact to Increase hPAAF Activation

A design of experiment approach further analyzed how each variable (microenvironmental stiffness, cell sex, serum sex, and serum age) and interactions among the variables tested influenced hPAAF activation. Microenvironmental stiffness had the greatest impact on hPAAF activation with a linear correlation between stiffness and activation (Figure 7A). Male hPAAFs, on average, were more responsive than female hPAAFs to increases in microenvironmental stiffness. There was a greater difference in activation of male hPAAFs between soft and stiffened conditions compared to female hPAAFs (Figure 7B). Age was also correlated with hPAAF activation. Age ≥ 50 serum showed greater activation levels than age < 50 serum even on soft conditions (Figure 7C). There was a larger difference in activation between soft and stiffened conditions in age < 50 serum than in age ≥ 50 serum. Next, hPAAF activation was correlated with individual sex hormone concentrations (estradiol, progesterone, and testosterone) quantified in each serum. A positive correlation between each sex hormone and increased activation was observed in stiffened conditions. Conversely, in soft conditions, sex hormones were negatively correlated with activation for both female and male hPAAFs (Figure 7D-F). Female cells showed higher activation levels on soft microenvironments than on stiffened microenvironments with low estradiol concentration. At low concentrations of progesterone or testosterone, differences in female hPAAF activation in response to microenvironmental stiffening were negligible; however, at higher concentrations, stiffening resulted in increased activation. Similarly, differences in male hPAAF activation levels between soft and stiffened conditions were smaller at lower sex hormone concentrations and grew with increasing concentrations of sex hormones.

**Figure 7.**
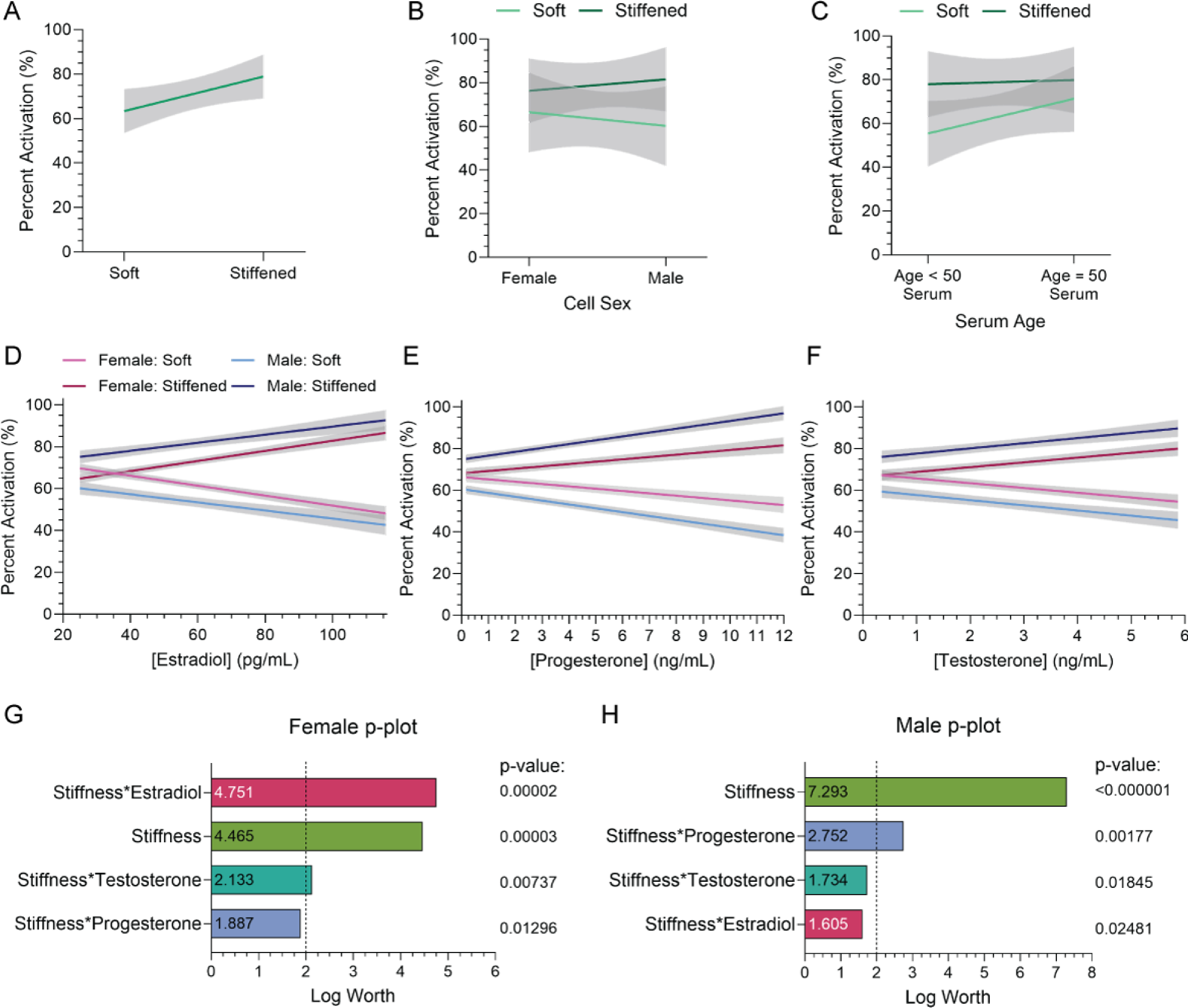
Interaction effect plots. Main effects plots revealed how variables A) Microenvironmental stiffness, B) cell sex and microenvironmental stiffness, and C) serum age and microenvironmental stiffness influenced hPAAF activation. Male hPAAFs tended to be more sensitive to changes in microenvironmental stiffness than female hPAAFs. Age < 50 human serum allowed for bigger changes in activation based on changes in microenvironmental stiffness, while age ≥ 50 serum tended to have higher activation levels regardless of microenvironmental stiffness. Interaction effect plots of D) estradiol concentration, E) progesterone concentration, and F) testosterone concentration in the serum with cell sex and stiffness. As sex hormone concentrations increased, activation on soft surfaces decreased, while activation of stiffened surfaces increased. Plots show mean ± 95% CI. P-plot of interaction effects on G) female hPAAFs and H) male hPAAFs. Female hPAAF activation was influenced the most by the compound effect of estradiol concentration and stiffness followed by stiffness, while male hPAAF activation was greatly influenced by stiffness followed by the compound effect of progesterone concentration and stiffness.

A combinatorial analysis of microenvironmental stiffness, estradiol concentration, progesterone concentration, and testosterone concentration effects on female or male hPAAF activation showed that microenvironmental stiffness was the factor most significantly driving male hPAAF activation. Interestingly, this was not true for female hPAAFs. Instead, the interaction between estradiol concentration and stiffness was the most influential driver of activation in female hPAAFs (Figure 7G-H). The magnitude of the effect of stiffness on male hPAAFs was 1.6 times greater than the magnitude of the stiffness effect on female hPAAFs. Microenvironmental stiffness was the factor with the second highest effect on female hPAAF activation while progesterone concentration combined with stiffness had the most influence on male hPAAF activation after stiffness alone.

## 4. DISCUSSION

Here, we quantified the differences in sex-specific hPAAF activation in response to increases in microenvironment stiffness while cultured in FBS, sex-matched or mismatched, age-separated human donor serum. Increased stiffness combined with older serum age led to the highest levels of fibroblast activation. Fibroblast activation and vessel stiffening are hallmarks of PAH pathogenesis [12, 25]. Prior studies have demonstrated the impact of substrate stiffness on all three of the pulmonary artery cell types. However, despite the known sex dimorphism in PAH [5–10], these studies did not distinguish between male and female cells and FBS (with undocumented hormonal milieu) was used as a supplement in the culture medium. Only a few studies have characterized the sex differences in fibroblast activation *in vitro*.

In this work, we found intrinsic sex-based differences in hPAAF activation in response to increases in microenvironmental stiffness and differences related to the species, sex, and age of the serum used to supplement the cell culture medium. When female and male hPAAFs were grown on dynamically stiffened hydrogels using FBS-supplemented media, female hPAAFs were less sensitive to increases in microenvironmental stiffness than male hPAAFs, with a significant increase in activation of male hPAAFs on stiffened surfaces compared to soft controls, but no change in activation of female hPAAFs with stiffening. However, female hPAAFs were more activated, even on soft conditions, than male hPAAFs. While we hypothesized that female hPAAF activation would likely be higher than activation of male hPAAF (since PAH predominately affects females over males [6–8]), the finding that female hPAAF were not responsive to stiffened conditions in FBS was not expected. A previous study investigating sex-specific human aortic valve leaflet fibroblasts that were cultured on soft and stiff substrates found that female cells were more sensitive to microenvironmental stiffness than male cells. In that study, female cells showed significant increases in activation on stiff hydrogels compared to soft hydrogel while grown in FBS, with no change in activation of the male cells on stiff hydrogels compared to soft hydrogels [15]. The differences in activation patterns found in our study and the published study may be due to inherent differences in responsiveness of these two cell types derived from different tissues.

The highest incidence of PAH is in females between the ages of 30-60 (average age = 36 ± 15 years old). Studies have shown that higher estradiol levels are associated with PAH development in both females and males [5–8]. Therefore, to study the influence of circulating sex hormones on hPAAF activation, we used human serum separated by sex and age. When assessing the effect of sex-matched, age-separated human serum on hPAAF activation, female hPAAFs responded to increases in microenvironmental stiffness when grown in pooled serum from female or male donors age < 50. We hypothesized that female hPAAFs would demonstrate increased activation when cultured with age < 50 female serum, as this serum had the highest pooled concentration of estradiol (115.7 pg/mL), and high estradiol levels are associated with increased PAH development. However, when cultured with serum from younger male donors where estradiol was low, female hPAAF were also responsive to microenvironmental stiffening and showed higher levels of activation on soft substrates. Interestingly, when grown with serum from donors ≥ 50 of either sex, hPAAF activation was higher on soft substrates, similar to the younger male serum, and there were no changes in activation with increases in stiffness, recapitulating what was seen when female hPAAF were grown in FBS. This result was interesting because estradiol concentration in the FBS was low, similar to the levels in the ≥ 50 female and male serum (Table 3), suggesting that estradiol may be a key regulator of activation levels in female hPAAF, especially on soft substrates.

Male hPAAFs cultured with age < 50 serum from either sex showed increased activation in response to stiffening, similar to levels measured when grown with FBS. When grown with age ≥ 50 serum, male hPAAFs demonstrated an increased activation on soft substrates and only showed responsiveness to stiffening when cultured with male ≥ 50 serum. The significant increases in activation of the male hPAAFs grown in the older male serum on the stiffened hydrogels could correlate to why males tend to have worse outcomes, as males tend to develop PAH at older ages than females do [6, 7, 26, 27].

We next used sex-mismatched serum to understand further the role of sex hormones versus innate sex differences in hPAAF activation. This experiment enabled the investigation of how sex-mismatch serum compositions and sex hormone concentrations impacted cellular activation in response to microenvironmental stiffening. Female hPAAFs showed increased levels of activation in serum from males of all ages and did not respond to microenvironmental stiffening. These findings contradict previous studies looking at the protective effects of testosterone within the pulmonary vasculature[7], as the male serum has much higher concentrations of testosterone than the female serum. These differences may be related to other differences in the serum composition. A study by Ramsey et al. found there are 77 differentially expressed analytes between female and male serum [19] and another study examining the impact of serum composition on fibroblast activation found that increased inflammatory cytokines in serum from patients undergoing aortic valve replacement increased aortic leaflet fibroblast activation [18].

Male hPAAFs showed similar activation levels on soft and stiff microenvironments when cultured with serum from young donors of either sex and with FBS. This suggests that the high levels of predominately female sex hormones (estradiol or progesterone) may allow cells to behave similarly under these different conditions. However, when male cells were grown in mismatched (female) age ≥ 50 serum, very high activation levels on both soft and stiffened conditions were measured, higher than when the male cells were grown in any other serum composition.

While studies have found that pulmonary vessels significantly stiffen with age [31], few studies have investigated how serum composition changes with age between the sexes outside of sex hormone concentrations. One study on serum biomarker levels between younger and older males found differences in nine types of cytokine levels between the age groups [32]. The results of our cytokine analysis supported the hypothesis that increased inflammatory signaling in older male serum could have initiated higher levels of activation; however, the same trend was not clear for female serum (Figure S3). Together, these data underscore the need to further identify components driving fibroblast activation in response to microenvironmental stiffening within human serum.

To validate the results observed in a commercially available male hPAAF cell line (2 years old), hPAAFs from three different male donors (24-30 years old) were cultured on dynamically stiffening hydrogels with age < 50 or age ≥ 50 male serum. Male PHBI donor fibroblasts exhibited levels of activation on soft and stiffened microenvironments comparable to the younger male hPAAF cell line. Male hPAAFs from the cell line showed a 1.43-fold change in activation while the average for the three donors was a 1.37-fold change in activation when supplemented with age < 50 male serum. In age ≥ 50 male serum, younger male hPAAFs exhibited a 1.39-fold change in activation and 1.21-fold change in activation in the PHBI donor fibroblasts. According to these results, the age of the male hPAAFs did not significantly influence the response to serum age or microenvironment stiffness.

To further analyze the effect of hormone concentration on female fibroblast response to increased microenvironmental stiffness, hPAAFs from three different female donors (28-36 years old) acquired from PHBI were cultured on dynamically stiffening hydrogels with age < 50, age ≥ 50, or age ≥ 50 plus estradiol female serum. These additional female PHBI donor fibroblasts showed activation levels on soft and stiffened hydrogels similar those measured in the commercially available female hPAAF cell line (11 years old). The commercially available female hPAAFs exhibited a 1.44-fold change in activation while the average of the three additional PHBI donor lines showed a 1.50-fold activation change in age < 50 female serum. A fold change in activation of 0.86 was measured in the female hPAAF cell line and 0.98 with female PHBI donor fibroblasts in age ≥ 50 female serum. The age of the female hPAAFs in this study did not significantly impact the response to serum age or microenvironmental stiffness.

When estradiol was added back to the age ≥ 50 female serum to match the estradiol concentration measured in age < 50 female serum, fibroblast activation increased on stiffened conditions compared to soft conditions and resembled the activation results in age < 50 female serum. A 1.36-fold change in activation was measured in these samples with a statistical difference between soft and stiffened. These results suggest that estradiol is an important regulator of female fibroblast activation even *in vitro*.

Analysis using a design of experiments approach demonstrated that microenvironmental stiffness and serum age were the only variables to independently have significant impact on fibroblast activation. Next, interaction effects were examined between the five factors studied here: substrate stiffness, cell sex, estradiol concentration, progesterone concentration, and testosterone concentration. The interaction between sex hormone concentrations and stiffness also significantly influenced fibroblast activation. As sex hormone concentrations increased, the fold change in activation between soft and stiffened conditions in both female and male cells also increased. This trend was consistent among all three sex hormones measured: estradiol, progesterone, and testosterone. Interaction analysis showed that at low concentrations of estradiol and progesterone (Figure 7D and E), female hPAAFs were not very responsive to stiffening and higher sex hormone concentrations increased female hPAAF sensitivity to microenvironmental stiffness. The analysis also confirmed our findings that male hPAAFs were sensitive to increases in microenvironmental stiffness and that sensitivity increased with higher sex hormone concentrations.

Microenvironmental stiffness was the factor in these experiments that influenced male hPAAF activation the most. Female hPAAFs, conversely, were most impacted by the compound effect of estradiol and stiffness (Fig 7G-H). Estrogens can increase prevalence of PAH but are also cardioprotective and protect against severe disease progression [3–5]. Interestingly, in this work, high levels of estradiol were correlated with lower levels of activation on soft surfaces while demonstrating increased activation on stiffened surfaces. These findings suggest that increased estradiol concentrations increase female hPAAF sensitivity to microenvironmental stiffness changes and therefore it may be possible to model the estrogen paradox observed in the PAH patient population *in vitro*.

Male hPAAFs were secondly impacted by the compound effects of progesterone and stiffness. These results were unexpected because progesterone is not typically considered a primary sex hormone in males, further highlighting the need to consider progesterone concentration when culturing male hPAAFs. The FBS evaluated here had the highest concentration of progesterone. Progesterone is the most dominant hormone present during pregnancy; therefore, it is unsurprising that FBS, isolated by blood collection from a fetal calf heart, has high concentrations of progesterone [3, 14]. However, combining male cells with FBS-supplemented medium is one of the most common modalities for investigating fibroblast activation *in vitro* [11, 12, 26–28]. Therefore, it is critical for researchers to report cell sex and serum hormone concentrations when performing activation assays *in vitro*.

Collectively, these results support the estrogen puzzle identified by other researchers in PAH and suggest it is possible to model it *in vitro*. Our study shows that higher levels of estradiol resulted in lower levels of activation on soft substrates replicating healthy tissue mechanics, leading to statistically significant increases in activation on stiffened surfaces, suggesting that estradiol increases female hPAAF sensitivity to changes in stiffness. Historic underrepresentation of female subjects in clinical trials, the failure of many studies to identify the sex of animals in *in vivo* studies, and a similar lack of information about tissue and cell sex in benchtop research have all perpetuated a long-standing disparity in healthcare outcomes for female patients. Thus far, the work in this study supports developments in the field of PAH investigating the biological differences of sex-specific pulmonary artery cell types and sex differences within circulating molecules. Future work will challenge hypotheses related to cellular activation in PAH in 3D to achieve a more physiologically relevant geometry of the pulmonary artery [21], which could include all three pulmonary artery cell types. Additional investigations could include incorporating cells and/or serum derived from PAH patients to better replicate PAH pathogenesis. A comprehensive analysis of other differential circulating factors in control versus PAH patient-derived serum would also further this investigation. It is important to continue studying sex differences to overcome healthcare disparities for all patients.

## 5. CONCLUSION

In this study, we measured differences in sex-specific hPAAF activation in response to increases in microenvironmental stiffness and sex-matched or mis-matched, age-separated human donor serum. Overall female hPAAFs tended to have increased levels of activation on soft and stiffened conditions, unless cultured in serum with relatively high concentrations of estradiol (e.g., younger female serum or older female serum with added estradiol). The results from this study suggest that higher sex-hormone concentrations increased female hPAAF sensitivity to increased microenvironmental stiffness. Male hPAAFs were more activated on stiffened microenvironments than on soft microenvironments regardless of the concentration of circulating sex hormones. This study also found that male hPAAFs were more sensitive to increases to microenvironmental stiffness, and sensitivity increased with higher sex hormone concentrations, especially in response to progesterone. Collectively, these results suggest that it is critical to intentionally design experiments considering cell sex and serum source as biological variables when conducting *in vitro* experimentation. Carefully controlling these aspects of cell culture will lead to improved models of the estrogen puzzle observed in PAH patients *in vitro*.

## Acknowledgement

This study was supported by the Ludeman Family Center for Women’s Health Research at the University of Colorado Anschutz Medical Campus to MCM and CMM; the Rose Community Foundation to MCM and CMM; the National Heart, Lung, and Blood Institute of the National Institutes of Health (NIH) under awards R01 HL153096 (CMM); the National Science Foundation under award 1941401 (CMM); and a Colorado Pulmonary Vascular Disease Research Award to CMM. The authors would like to thank the Colorado Clinical Translational Science Institute for performing sex hormone concentration analysis supported by CTSA Grant UL1 TR002535, KL2 TR002534, & TL1 TR002533. Cells were provided by the Pulmonary Hypertension Breakthrough Initiative (PHBI). Funding for the PHBI is provided under an NHLBI R24 grant, #R24HL123767, and by the Cardiovascular Medical Research and Education Fund (CMREF).

## Data Availability

The data that support the findings of this study are openly available at the following URL/DOI: https://doi.org/10.17632/gs44hgz895.3.

