## Supplementary Material for "Dynamically stiffening biomaterials reveal age- and sex-specific differences in pulmonary arterial adventitial fibroblast activation"

**NMR Results**

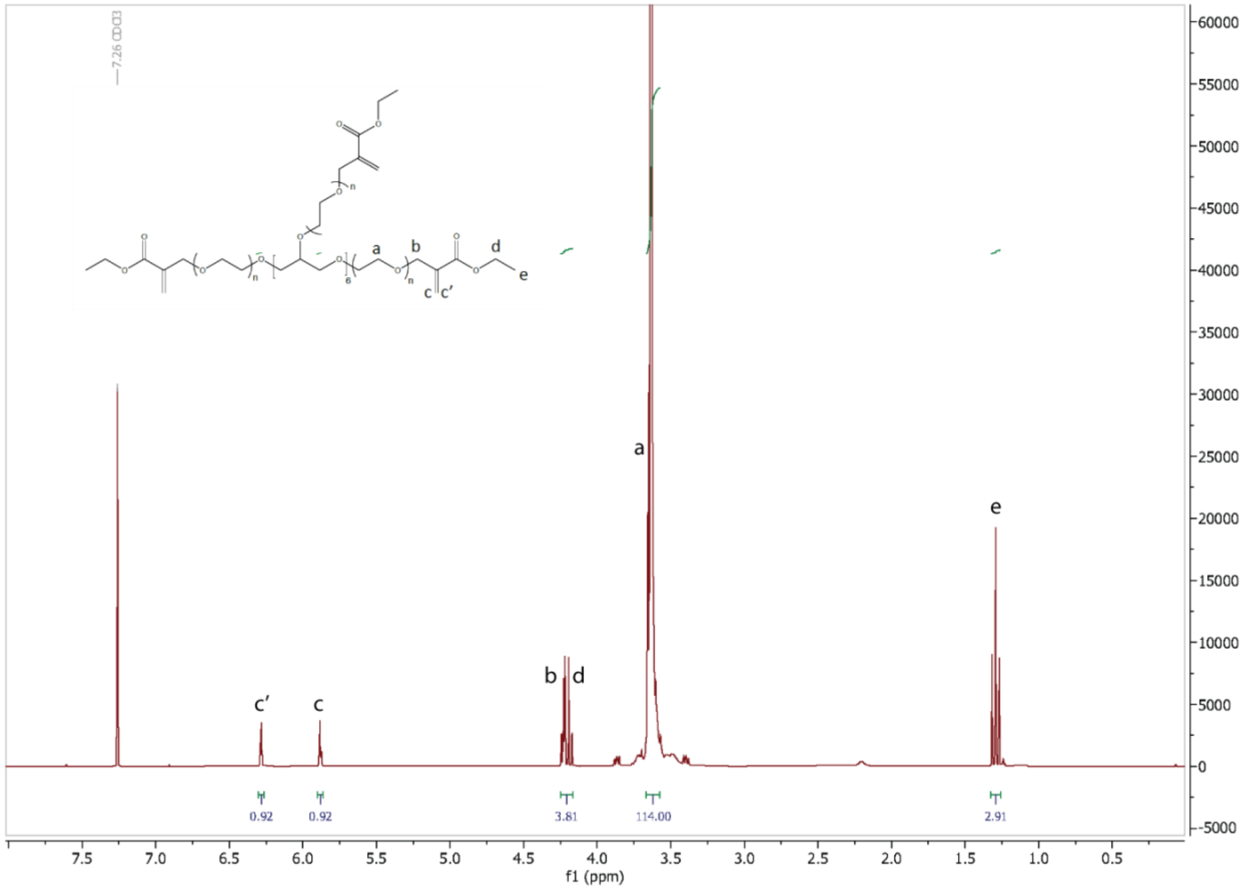


**Figure S1.** PEGαMA 1H NMR (300 MHz, CDCl3): δ (ppm) 1.36 (t, 3H, CH3–), 3.71 (s, 114H, PEG CH2-CH2), 4.29 (t, s, 4H, –CH2–C(O)–O–O, –O–CH2–C(=CH2)–), 5.93 (q, 1H, –C=CH2), 6.34 (q, 1H, –C=CH2). End group functionalization of the final PEGαMA polymer was greater than 92% by comparison of the αMA alkene end group to the PEG backbone.

### Soft Hydrogel Activation Results

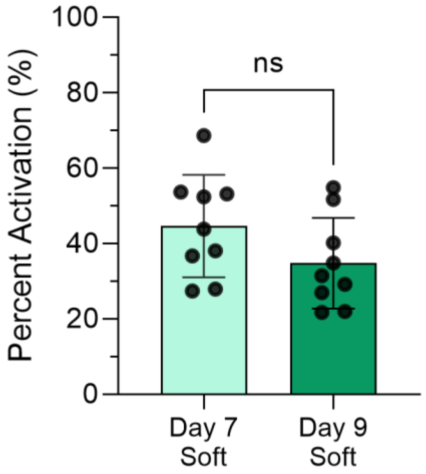


**Figure S2.** Activation of male hPAAFs grown in FBS on soft hydrogels was not significantly different when comparing cells collected on Day 7 to Day 9. Columns represent mean ± SD, n = 3, Mann-Whitney.

### Western Blot Methods

Protein was isolated from samples using TRI Reagent (Molecular Research Center) with 20 mg/mL of tRNA from S. cerevisiae (Sigma) using manufacture's protocol. Protein was resuspended in 8 M urea (Sigma), 2.5 M thiourea (Sigma), 4% CHAPS (Sigma), 2 mM 4-(2-hydroxyethyl)-1-piperazineethanesulfonic acid (EDTA, Sigma), 10 mM DTT, 1% tributylphosphine (BioRad), and 1% protease inhibitors (Sigma). Protein concentration was determined using a modified protein assay (BioRad) and prepared in Laemmli sample buffer (BioRad). SDS-PAGE was performed using 10% acrylamide gel (30% acrylamide/0.8 Bis (BioRad), 1.5M Tris (Fisher Scientific), 20% sodium dodecyl-sulfate (SDS, Sigma), 10% ammonium persulfate (BioRad), tetramethylethylenediamine (BioRad), and proteins were transferred to PVDF membrane (BioRad). Membranes were blocked with 5% BSA in Tris-buffered saline with 0.1% Tween 20 ((Sigma), TBST). Membranes were incubated in anti-actin α-smooth muscle (Sigma, A2547) at 1:500 dilution overnight at 4°C. Membranes were washed three times with TBST before incubation with secondary antibody (goat anti-mouse 1:25,000; Sigma, A2304) for 1 hour at room temperature. Membranes were washed three times with TBST before being developed in ECL (Thermo) and imaged. Blots were quantified using volume analysis densitometry in ImageJ.

**Western Blot Results**

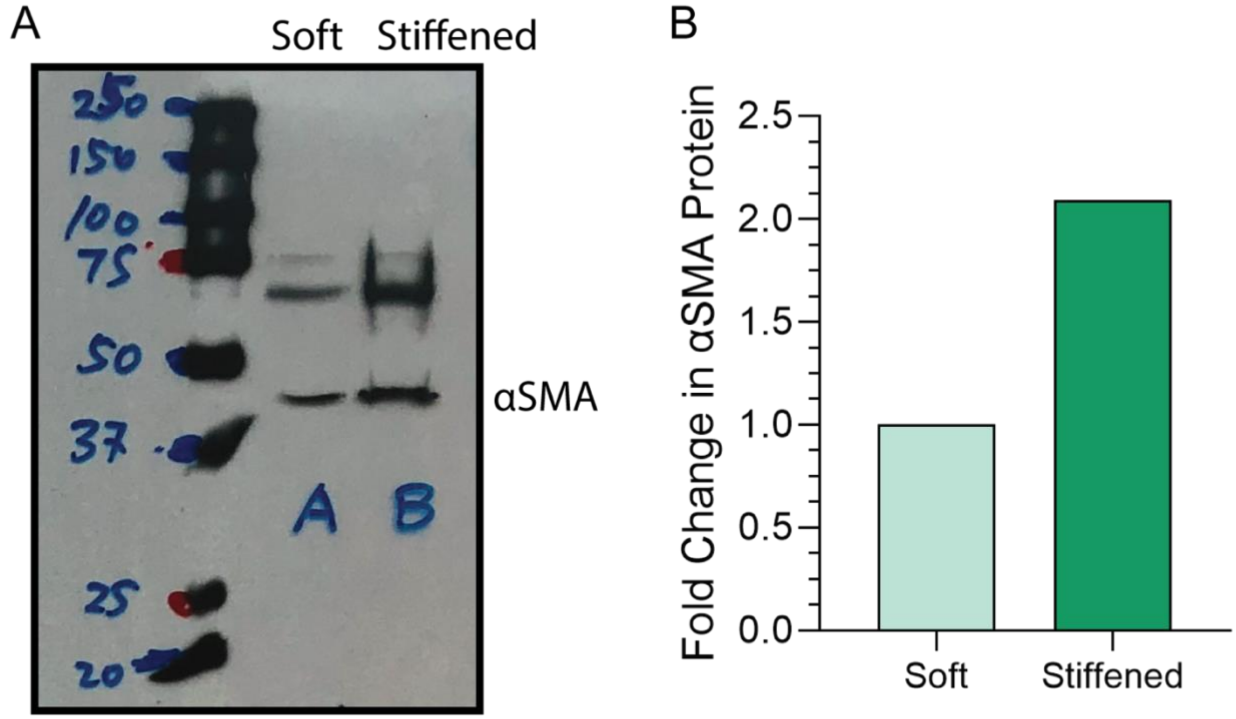


**Figure S3.** Western blot of αSMA protein expression in male hPAAFs supplemented with FBS on soft verses stiffened hydrogels (n = 3).

### Serum Cytokine Array Methods

An antibody-based array for a panel of human cytokines (Abcam ab133997) was used according to the manufacturer's protocol. Briefly, human serum pooled by sex and age (N=3 technical replicates) were mixed 1:1 with blocking buffer and applied to pre-blocked membranes for an overnight incubation at 4°C. Samples were washed and sequentially incubated in biotinylated antibodies and streptavidin-HRP. Each incubation ran overnight at 4°C. After a final wash, membranes were developed and imaged on a C-DiGit Blot Scanner (Li-Cor). Blots were quantified using volume analysis densitometry in ImageJ, and the results were normalized to positive controls after subtraction of negative controls.

### Serum Cytokine Array Results

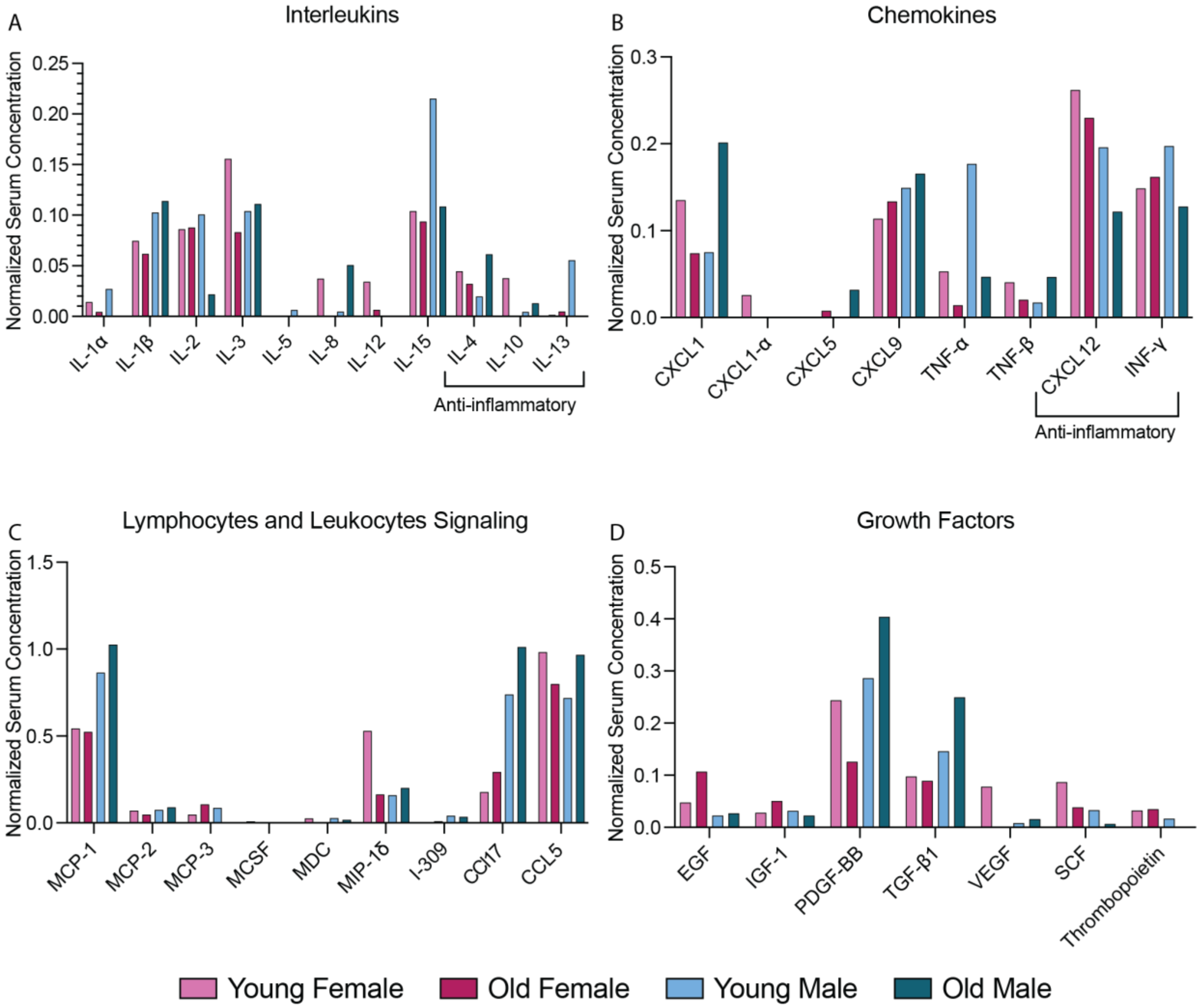


**Figure S4.** Several circulating biochemical cues including A) interleukins, B) chemokines, C) lymphocyte and leukocyte signaling molecules, and D) growth factors were detected within human serum samples pooled by sex and age.

**Vehicle Control Results**

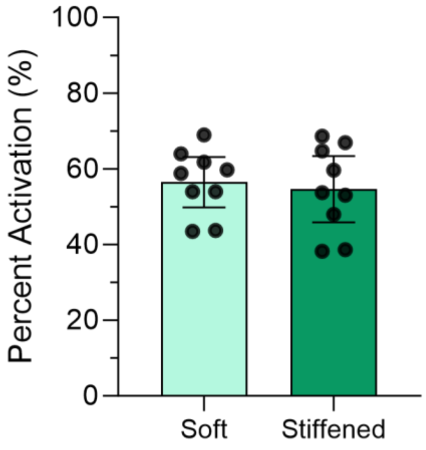


**Figure S5.** Female donor hPAAF activation in age ≥ 50 female human serum with ethanol vehicle control cultured on soft or stiffened hydrogels was not statistically different from activation in age ≥ 50 female human serum alone. Columns represent mean ± SD, n = 3, Mann-Whitney.
